# HSeeker: an algorithm for systematic H-DNA sequence identification

**DOI:** 10.64898/2026.07.10.737678

**Authors:** Kimonas Provatas, Guliang Wang, Nikol Chantzi, Archit Patil, Imee MA Del Mundo, Candace SY Chan, Ilias Georgakopoulos-Soares, Karen M Vasquez

## Abstract

H-DNA is a naturally occurring intramolecular DNA triplex structure formed by Hoogsteen hydrogen bonds at homopurine-homopyrimidine mirror repeats and has functional roles in gene regulation, genome instability, and human disease. The existing H-DNA detection tools capture only a subset of H-DNA sequences, often missing relevant sequence features or failing to assess key aspects of structural stability. To address this gap, we present “HSeeker”, a state-of-the-art computational tool that compiles a three-part algorithm to identify and score potential H-DNA-forming sequences. Using a center-outward search algorithm approach, HSeeker evaluates candidate hinge positions and spacer lengths while allowing configurable mirror mismatches and purine-pyrimidine composition thresholds. The greedy overlap removal phase resolves overlapping candidates by retaining the longest and most compact motif within each overlapping region. Finally, the thermodynamic stability scoring algorithm evaluates the candidate motifs using an experimentally informed scoring model that incorporates Hoogsteen G-G and A-A bonds, consecutive-pair stacking, and imposes penalties for mismatches and disrupted stacks. The scoring procedure also optimizes motif boundaries by trimming weak terminal positions and reassigning unstable arm positions to the spacer. HSeeker reports genomic coordinates, sequence information, pairing and stacking components, and an overall stability score. HSeeker is also user-friendly, available as a Python package and as a web application, supporting configurable, high-throughput analysis and exportability of predicted H-DNA motifs. HSeeker provides an accessible and reproducible framework for investigating the distribution and potential stability of H-DNA-forming sequences across genomic datasets.

## Introduction

H-DNA is an alternative (i.e., non-B) DNA structure that forms at homopurine-homopyrimidine mirror-repeat sequences. H-DNA is formed by the folding back of one of the repeat arms and Hoogsteen hydrogen bonding to the purine-rich strand of the underlying duplex through the major groove, generating a triple-stranded structure, and leaving the complementary strand partially single-stranded ^1,2^. When a pyrimidine-rich sequence serves as the third strand, it adopts a parallel orientation relative to the purine-rich strand of the underlying duplex, stabilized by canonical Hoogsteen hydrogen bonds. Notably, the formation of this parallel motif typically requires cytosine protonation to generate C+.G-C triads. This pH dependence makes the structure biologically less likely to form under standard intracellular conditions. Conversely, antiparallel H-DNA forms when a purine-rich strand folds back to interact via reverse-Hoogsteen hydrogen bonds with the purine-rich side of the duplex in an antiparallel orientation. The antiparallel conformation maintains stability at neutral physiological pH compared to its parallel counterpart, rendering it a more viable structural motif *in vivo*.

H-DNA structures can arise from and also modulate essential DNA metabolic processes with important roles in genome function and stability^3^. For example, H-DNA structures can interfere with replication, transcription, DNA repair^4–6^, often by acting as an impediment to l polymerases^3,7,8^. With the exposed single-strand, it can also accumulate more damage and repair errors, resulting in increased mutagenesis^9–11^. Consistent with this model, naturally occurring H-DNA-forming sequences have been shown to be intrinsically mutagenic in mammalian cells and in mice^12,13^, and often overlap with human disease-causing mutations^14–16^, supporting the idea that structure-forming DNA sequences are an endogenous source of genome instability. Thus, H-DNA motifs are considered candidate mutational hotspots, particularly in genomic regions where replication or transcription imposes negative supercoiling, which favors the formation of alternative DNA structures.

One of the best-studied examples of H-DNA-associated instability occurs at the human *c-MYC* locus^17–19^. The *c-MYC* promoter contains sequences capable of adopting a variety of non-B DNA conformations, including H-DNA, and these regions overlap with sites implicated in transcriptional regulation and genetic instability^19^. Because *c-MYC* is a key oncogene whose dysregulation contributes to multiple cancer types, the presence of H-DNA-forming sequences at this locus has provided an exemplary case study for understanding how H-DNA can influence genomic instability and gene regulation associated with cancer. More broadly, H-DNA and related triplex-forming motifs have been linked to chromosomal fragility, rearrangement-prone regions, and disease-associated instability, suggesting that systematic annotation of H-DNA-forming potential could improve our understanding of genome organization, mutational processes, and disease etiology.

Despite this biological relevance, to fully understand the impact of H-DNA on cellular functions, a better understanding of its distribution in genomic sequences is warranted. While we know that potential H-DNA-forming sequences are abundant in mammalian genomes and are possibly enriched in promoters and introns, computational identification and annotation of H-DNA-forming sequences remain less developed than for several other non-B DNA structures. A comprehensive mapping across different genomes is also lacking. Early approaches searched for homopurine-homopyrimidine mirror repeats using relatively simple sequence rules, while later efforts introduced more flexible models. For example, Schroth and Ho proposed sequence-based criteria for identifying potential H-DNA-forming motifs^20^, and Lexa *et al*. (2011) subsequently developed a dynamic programming algorithm for detecting triplex-forming sequences that allowed imperfect matches rather than restricting searches to perfect mirror repeats^21^. These advances were important because experimentally validated H-DNA-forming sequences may contain interruptions, mismatches, or variable spacer lengths, features that are missed by overly strict pattern-matching approaches.

Few computational tools are currently available to detect H-DNA-forming sequences; however, these existing resources have important limitations for systematic H-DNA discovery. For example, nBMST provides a general interface for searching multiple classes of non-B DNA motifs, but its implementation only includes mirror repeats of at least 90% purines or pyrimidines, without tolerating any mismatch in mirror symmetry ^22,23^. NeSSie is a C/C++ library and tool optimized for detecting the mirror repeats characteristic of H-DNA, utilizing an optimal global alignment based on the Needleman-Wunsch algorithm^24^, and allowing gaps, mismatches, or asymmetric loops to be present. Triplexator^25^ and 3plex/3plex Web^26^ were both originally developed for RNA:DNA intramolecular H-DNA and intermolecular Triplex Target Site prediction, which is related but distinctly different from intramolecular H-DNA formation *in vivo*, and they did not provide a dedicated genome-scale annotation framework for intramolecular H-DNA-forming mirror repeats^25^. 3plex is the only pipeline that integrates structural rules with thermodynamic modeling. The algorithm first evaluates the canonical rules for parallel and antiparallel Hoogsteen base-pairing, followed by a procedure to evaluate the thermal stability profiles derived from DNA denaturation experiments. It allows calculating an empirical affinity score for a single-stranded DNA or RNA, rather than relying solely on strict sequence patterns^26^. However, since it was designed to search for intermolecular triplexes, the thermal parameters were derived from single-stranded DNA or RNA, and the existing length and the length of the mirror-symmetric center were not taken into consideration. As a result, there remains a need for dedicated software that can systematically identify naturally occurring intramolecular H-DNA motifs while accounting for biologically relevant sequence features such as mirror symmetry, homopurine-homopyrimidine composition, spacer length, and imperfect triplex-forming potential.

Here, we present “HSeeker”, a computational framework for the systematic identification of anti-parallel H-DNA-forming sequences from genomic DNA. By integrating sequence composition, mirror-repeat architecture, and tolerance for imperfect triplex-forming patterns, HSeeker enables scalable annotation of highly probable candidate H-DNA loci across genomes. This framework provides a foundation for studying the distribution, evolution, and mutational consequences of H-DNA-forming sequences, and for integrating H-DNA annotations into broader analyses of non-B DNA, genome instability, and disease-associated variation.

## Methods

### H-DNA candidate sequence discovery

HSeeker searches for antiparallel H-DNA that contains reverse Hoogsteen hydrogen bonds (e.g., G·G-C and A·A-T triplets) for their occurrence in neutral physiological conditions (**Figure 1**). It employs a center-outward heuristic to detect potential H-DNA-forming mirror repeats in genomic sequences, augmented with an exact purity-RMQ prefilter that eliminates center-spacer pairs whose right arm cannot satisfy the composition constraint. For each potential hinge center *c* and spacer length *s*, the right-arm start is *r*_0_ = *c* + *s* + 1. Before performing character-by-character extension, HSeeker computes the maximum reachable arm length *K*_max_ = min(*L*_max_, *c +* 1, *n* − *r*_0_, nextBad(*r*_0_) − *r*_0_),, where nextBad(*r*_0_) is the first non-ACGT position at or after *r*_0_. It then uses prefix-count arrays *G* (*i*) and *C* (*i*) for GA-rich and CT-rich right-arm bases and defines 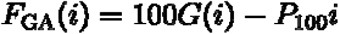 and 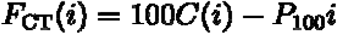, where *P*_100_ = [100*P* + 0.5] is the integer-scaled purity threshold. A GA-rich right-arm prefix of some length *k* ∈ [*L*_min_, *K*_max_] can meet the required purity threshold exactly when 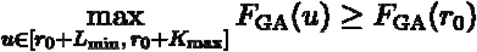, and the CT-rich case is tested analogously using *F*_CT_. These two range-maximum queries are a deterministic feasibility check: if neither inequality holds, no extension length for that center-spacer pair can pass the original purity rule, and the pair is skipped without changing output semantics. All remaining pairs are processed by the original center-outward extension logic: the algorithm compares left and right bases, increments the mismatch counter on differences, tracks GA/CT composition from the right arm, evaluates mirror identity and purity using integer-scaled arithmetic, and retains only the longest valid arm for each (*c, s*) pair. Extension terminates when the mismatch budget is exhausted or a non-ACGT right-arm base is encountered, as described in **Algorithm 1**. Following the scanning phase, overlapping hits are resolved by a greedy linear sweep over hits sorted by genomic start: the hit with the longest arm survives each overlap group, with ties broken by shorter spacer length. The candidate enumeration step is described in **Algorithm 2**.

**Figure 1:**
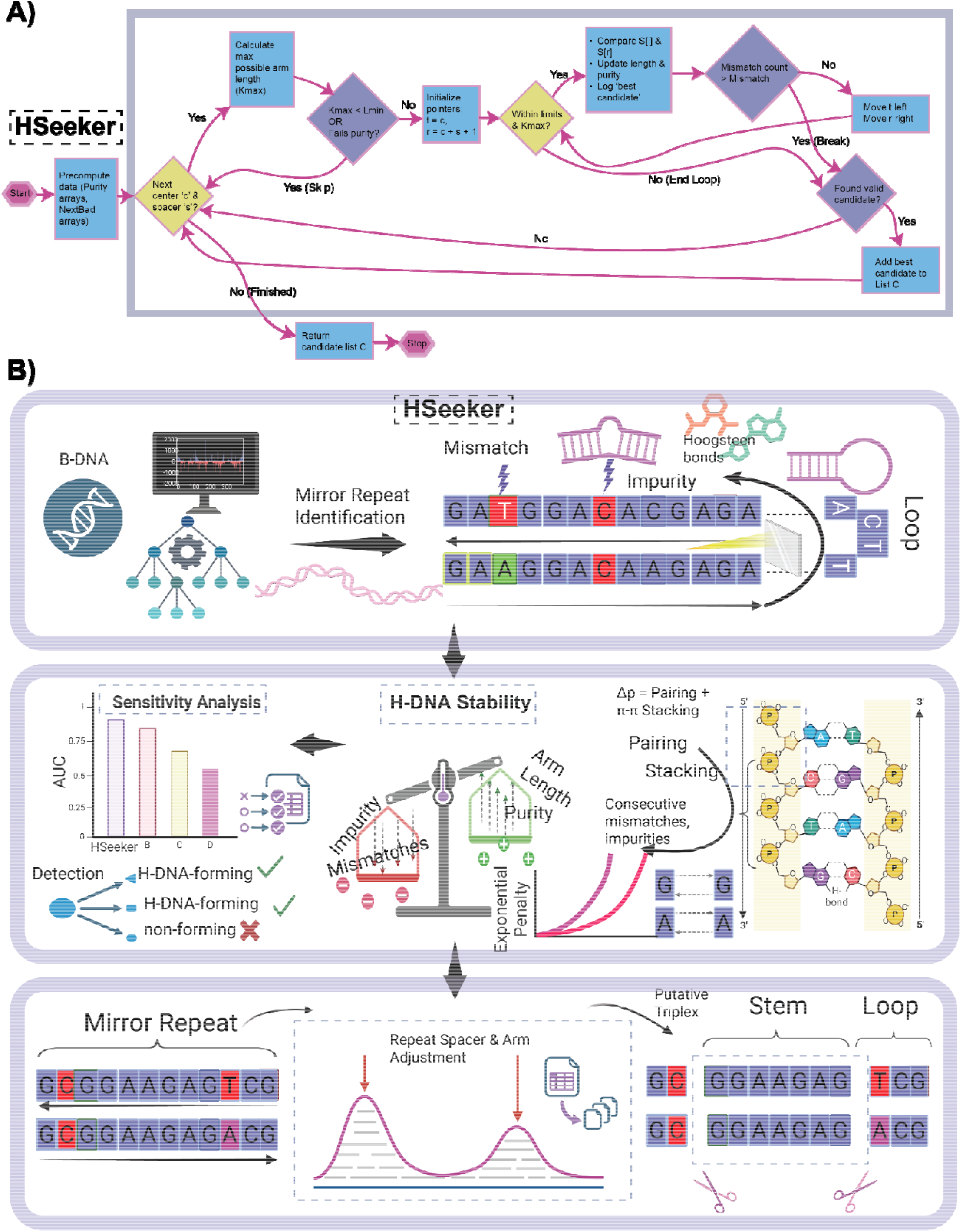
Computational workflow for the identification and scoring of putative H-DNA-forming mirror repeats. **A)** HSeeker candidate-identification workflow. **B)** Starting from a sequence input, the pipeline identifies homopurine/homopyrimidine mirror repeats and detects sequence features that may disrupt triplex formation, including mismatches, impurities, and loop-forming segments. Identified candidates are evaluated using an H-DNA thermostability score that integrates sequence composition and structural determinants (e.g., mismatches/impurities, repeat purity, arm length, base pairing, and base stacking) to estimate the propensity of each sequence to form a stable H-DNA/triplex structure. Candidate mirror repeats are further refined by adjustment of repeat spacer and arm boundaries to define the final putative triplex architecture, consisting of a triplex-forming stem and an intervening loop. Together, this framework enables systematic detection and prioritization of sequences with the potential to adopt H-DNA structures.

### Sequence composition filters

Prior to stability scoring, we applied two empirical filters based on experimentally derived criteria for H-DNA formation. First, we filtered out all hits where the mirror repeat arm had an AT-content ≥80% ^20^, as sequences with very high AT-richness are known to destabilize triplex formation due to geometric incompatibility of Hoogsteen base triplets and reduced stability of A·A-T triads. Second, following stability scoring, we excluded any putative triplex that consisted of a homopolymeric run (poly-A, poly-G, poly-C, or poly-T), as these do not form stable inter-strand triplex interactions despite satisfying homopurine/homopyrimidine requirements.

### Stability score

The stability scoring was based on the exact interaction energy changes for each Hoogsteen base pair within a DNA triad provided by Park *et al*. (2021)^27^, isolated and calculated by simulating the triplex structures without the constraints of the phosphate backbone. Therein, the interaction energy of a G·G reverse Hoogsteen hydrogen bond in a G·G-C triplet was 7.63 kcal/mol, and that of an A·A reverse Hoogsteen hydrogen bond in an A·A-T triplet was 3.83 kcal/mol. In addition, Park *et al*. (2021) calculated the interaction energies of different combinations of stacking within the 3rd strand in an H-DNA structure, which ranged from 2.15 to 6.65 kcal/mol. To simplify the searching and scoring, we took an average of 5.0 kcal/mol of stacking energy penalty if a mismatched base exists in a third strand. Taken together, the HSeeker stability scoring was calculated as follows:

Let 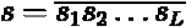 denote an H-DNA-forming sequence, where *L* > 0 is the arm length. The stability is defined as the total sum of the energy of pairing and stacking interactions of the H-DNA-forming sequence. In particular, we used a greedy algorithm to identify the subsequence of a mirror repeat with the maximum stability, assessed using a prefix and suffix maximization algorithm. The stability is dependent entirely on the mirror repeat arm and not on the spacer length. However, H-DNA is thermodynamically unfavourable with very long spacer lengths; thus, we imposed a constraint on the total spacer length that we consider at a maximum of 20 bp. In particular, let *m* > 0 denote the spacer length. We require that *m* ≤ min (20, *L*).

For each mirror repeat, we consider two types of intramolecular forces: pairing and π-π stacking energies. Again, we assume that only the anti-parallel formation is likely to form, considering neutral pH conditions. Given a purine-dominant sequence, we consider the values in **Supplementary Table 1**. Given the score table, we can now attempt to extract the subsequence anchored at the spacer with the maximum consecutive score.

Each candidate hit produced by the purity-prefiltered center-outward enumeration and surviving overlap removal is subsequently evaluated for thermodynamic stability. The scorer receives the full motif sequence *s* = *s*_1_ · *s*_3_ · *s*_2_ ·, composed of left arm *s*_1_, spacer *s*_3_, and right arm *s*_2_, together with the detected arm length *L*_arm_. If *s*_1_ is CT-rich (i.e.,|*CT*(*s*_1_)| > |*GA*(*s*_1_)|), the full sequence is reverse-complemented so that scoring always proceeds on the purine-rich strand. The right arm is then reversed to yield 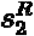, aligning position *j* of 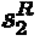 with position *j* of *s*_1_ for direct positional Hoogsteen comparison. Non-canonical 5’ positions — those where neither 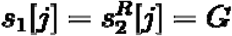 nor 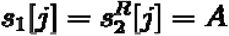 holds — are stripped from both arms before scoring. A per-position pairing score is then assigned as *P* [*j*] = 7.6 for G--G pairs,*P*[*j*] = 3.83 for A--A pairs, and *P*[*j*] = − (7.6 + 3.83) / 2 for any mismatch in the body of mirror repeat or impurity, Furthermore, a stacking bonus of +5 is awarded at each position where the two adjacent positions are both canonical. In particular, we penalized the π-π stacking mismatches or impurities using the formula − *δ*^*n*+1^/*d*, where *n* counts the length of the immediately preceding non-canonical run. Let *f* (*s*) denote the scoring array considering pairing and stacking energies. Since consecutive mismatches or impurities would destabilize the H-DNA structure, we modeled the penalties in π-π stacking as exponential. The putative H-DNA-forming sequence is defined as the subsequence of *s* [*L*^∗^: *R*^∗^], such that (*L*^∗^: *R*^∗^) maximizes the stability of *ℰ*, defined as follows:

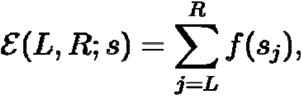

subject to the following constraints:

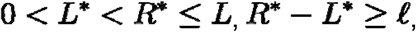

and

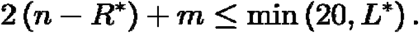

Thus,

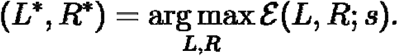

The procedure yields a stacking score, a pairing score, their sum as the total score, and a boundary-optimised putative triplex in *s*_*l*_[*s*_spacer_]*s*_*r*_ notation. The scoring procedure is described in **Algorithm 3**.

### Sensitivity analysis

To assess the ability of HSeeker to detect H-DNA-forming sequences, we conducted a sensitivity analysis using a benchmark set of 79 experimentally validated sequences: 70 known to form H-DNA and 9 negative controls. Positive controls were selected based on strong experimental evidence of H-DNA formation, including nucleotide-level structural fine-mapping via chemical modification assays in a plasmid context (**Supplementary Table 1**).

*Direct sequence-level validation*. We first benchmarked HSeeker against two existing tools, Triplex and Triplexator, running all three independently on the 79 benchmark sequences. HSeeker was run with its default parameters: *hseeker -seq hdna_benchmark_injected*.*fna -minrep 8 -maxspacer 10 -mismatch 0*.*15 -purity 0*.*9 -maxrep 1000 -out hdna_benchmark*

Triplex (v1.50.0) ^21^ was run through its R interface (triplex.search) using a minimum score of 15, length bounds of 8–50 bp, and prokaryotic EVD parameters: *triplex*.*search(fasta, min_score=15, min_len=8, max_len=50, seq_type=“prokaryotic”)*

Triplexator (v1.3.2) ^25^ was run with a 10 bp minimum length (the tool’s lower limit), no maximum length cap, a 15% error rate (matched to HSeeker’s mismatch tolerance), and the repeat filter disabled to avoid excluding valid H-DNA motifs: *triplexator -ds <fasta> -l 10 -L -1 -e 15 -fr off -o <out>*

A sequence was scored as positive if any tool reported at least one candidate H-DNA motif. To test whether HSeeker’s stability score could distinguish forming from non-forming sequences beyond simple detection, we identified an optimal classification threshold using Youden’s J statistic, which balances true-positive and true-negative rates. We also generated a ROC curve by applying the stability score across all detected hits.

#### Genomic context validation

We next tested whether each tool could recover the benchmark sequences when embedded in a realistic genomic background. We downloaded the *E. coli* K-12 genome from NCBI and randomly inserted each of the 79 sequences into it, enforcing a minimum 500 bp spacing between insertions (fixed random seed, seed=42). HSeeker was run on the resulting genome using the same parameters as above. A detected hit was counted as a true match if its overlap with the insertion site exceeded 80% of either the hit length or the insertion length, a threshold chosen to accommodate cases where tools extend hits into flanking sequence or detect only the triplex core within a longer insertion.

For both the direct and genomic-context analyses, performance was evaluated using AUC-ROC and the Matthews Correlation Coefficient (MCC), the latter chosen to account for the substantial class imbalance in the dataset (∼7.8:1 ratio of H-DNA-forming to non-forming sequences).

##### Algorithm 1

HSeeker Phase 1: Candidate Enumeration with Purity RMQ Prefilter

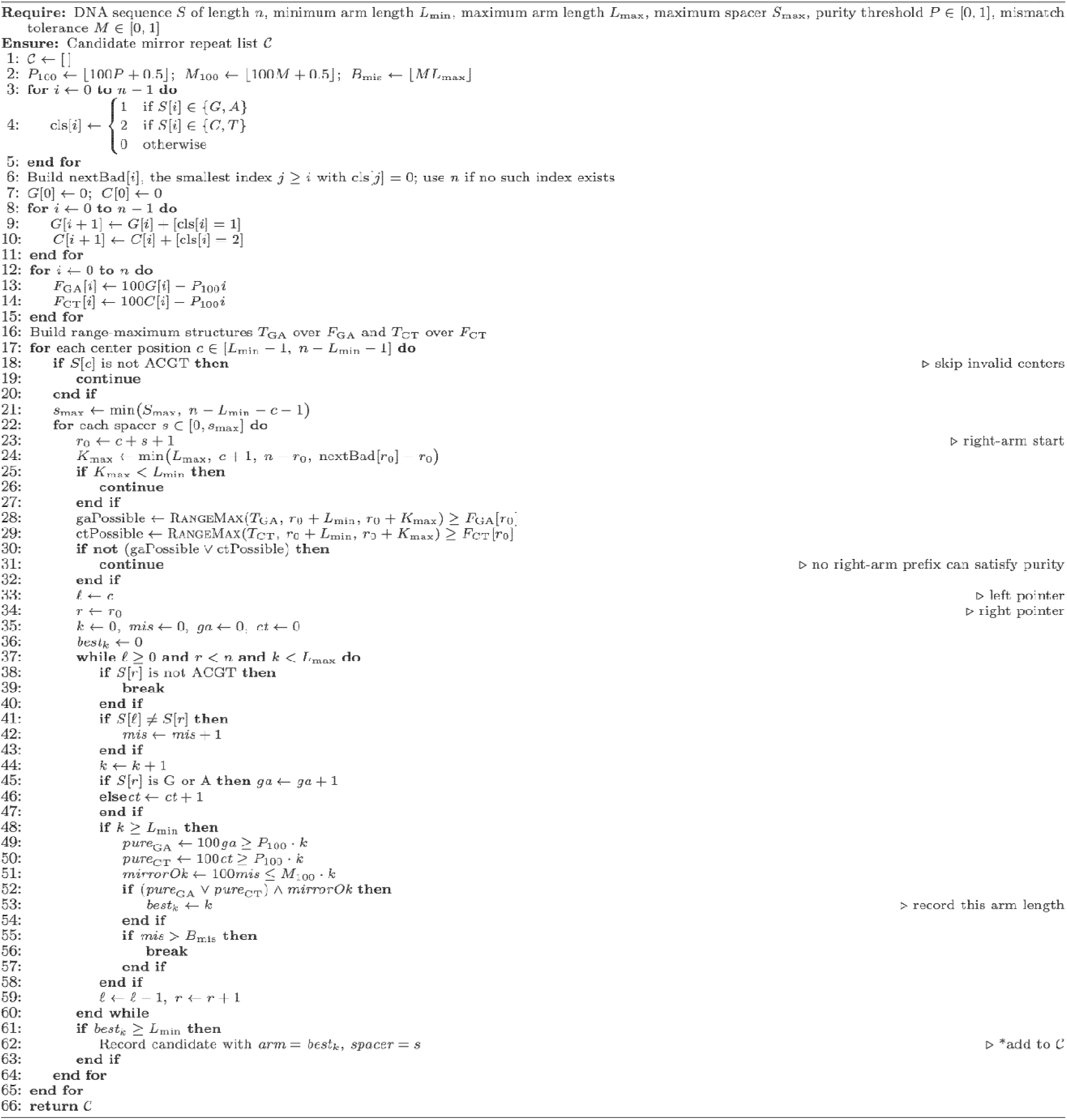

**Algorithm 1: Center-outward mirror repeat candidate enumeration with purity RMQ prefilter**

HSeeker first constructs right-arm composition prefix arrays and range-maximum query structures that test whether any admissible right-arm prefix can satisfy the GA-rich or CT-rich purity threshold. For each valid center position and spacer length, this prefilter computes the maximum reachable arm length and skips the center-spacer pair if no prefix in the admissible length interval can pass purity. Candidate pairs that pass the prefilter are then extended outward exactly as in the original scanner: the left and right pointers move simultaneously, mirror mismatches and right-arm compositional purity are tracked, and an arm is recorded only when both mirror identity and purity meet their thresholds. Only the longest valid arm for each center-spacer pair is retained.

##### Algorithm 2

HSeeker Phase 2: Overlap Removal

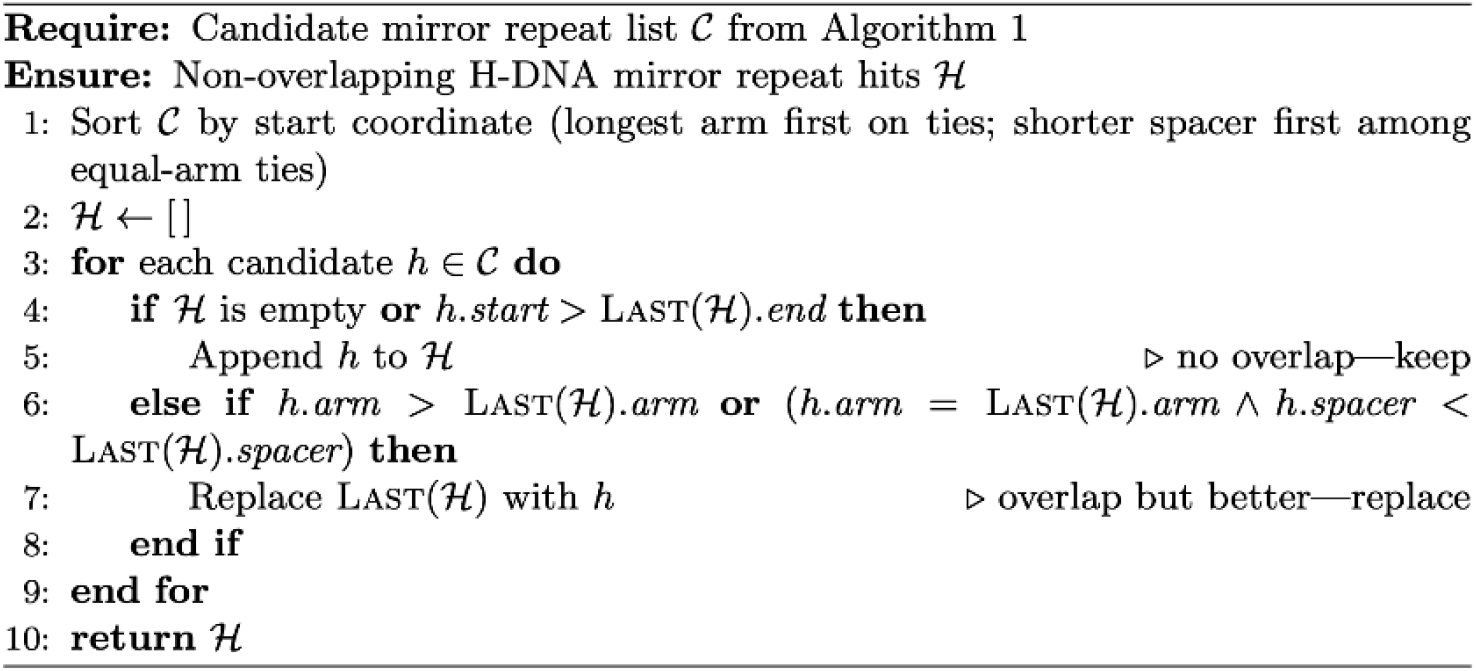

**Algorithm 2: Greedy overlap removal**

All hits are sorted by the genomic start coordinate, with longer arms prioritised on ties. A single linear sweep keeps non-overlapping hits and, among overlapping hits, retains the one with the longest arm (or equal arm and shorter spacer). The result is a compacted list with no overlapping hits.

### Complexity analysis

HSeeker processes each input sequence through candidate enumeration, overlap removal, and thermodynamic scoring, with costs governed by the sequence length *n*, maximum spacer *S*_max_, maximum arm length *L*_max_, and minimum arm length *L*_min_. With the purity-RMQ optimization enabled, candidate enumeration first performs a linear preprocessing pass over the sequence to classify bases, build the right-arm invalid-position array nextBad, construct GA/CT prefix-count arrays, and derive the transformed purity signals 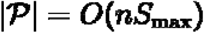 and 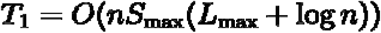 . Building the range-maximum structures over these signals costs *O*(*n*) time and *O*(*n*) memory for the segment-tree implementation. The scanner still considers *O*(*n*) center positions and *O*(*S*_max_) spacer lengths per center. For each (*c, s*) pair, it computes *r*_0_ = *c* + *s* + 1 and *K*_max_(*c, s*) = min(*L*_max_, *c* + 1, *n* − *r*_0_, nextBad(*r*_0_) − *r*_0_), then performs two range-maximum queries to test whether any right-arm prefix length *k* ∈ [*L*_min_, *K*_max_] can satisfy the GA-rich or CT-rich purity condition. These feasibility checks cost *O*(*log n*) per (*c, s*) pair in the current segment-tree implementation and can skip the center-outward extension entirely when no valid-purity right arm exists. If a pair passes the prefilter, then the original extension loop runs for at most *K*_max_(*c, s*) ≤ *L*_max_ steps, tracking mirror mismatches and retaining the largest valid arm. Thus the Phase 1 cost is 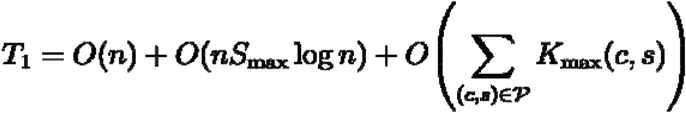, where *P* is the subset of center-spacer pairs that pass the purity prefilter. In the worst case, all pairs may be purity-feasible, so |*P*| = *O*(*nS*_max_) and *T*_1_ = *O*(*nS*_max_(*L*_max_ + log*n*)), which simplifies to for practical settings where *L*_max_ ≫ log*n*. The optimization therefore preserves the original worst-case asymptotic bound but substantially reduces practical work by eliminating inner-loop extensions for purity-impossible pairs. Candidate enumeration produces at most *H* ≤ *nS*_max_ raw candidates, at most one per center-spacer pair; overlap removal sorts them in *O*(*H* log *H* ) time and performs a linear sweep in *O*(*H*), giving *T*_2_ = *O*(*H* log *H*) = *O*(*nS*_max_ log(*nS*_max_)). After overlap removal, surviving hits are pairwise non-overlapping, and since each spans at least 2*L*_min_ bases, the number of scored hits is *H*′ ≤ ⌊*n* / (2*L*_min_) ⌋ = *O*(*n*/*L*_min_). Stability scoring applies a constant number of linear passes over each hit’s arms and adjusted spacer boundaries, each bounded by *O*(*L*_max_), so *T*_3_ = *O*(*H*′ *L*_max_) = *O*(*nL*_max_/*L*_min_). The full optimized pipeline is therefore 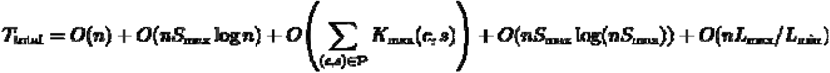, with worst-case simplification *T*_total_ = *O*(*nS*_max_*L*_max_). Thus, purity RMQ does not change the theoretical worst-case class, but it changes the practical cost driver from extending every center-spacer pair to extending only the subset that can possibly satisfy the right-arm purity threshold.

#### Algorithm 3

HSeeker Phase 3: Stability Scor

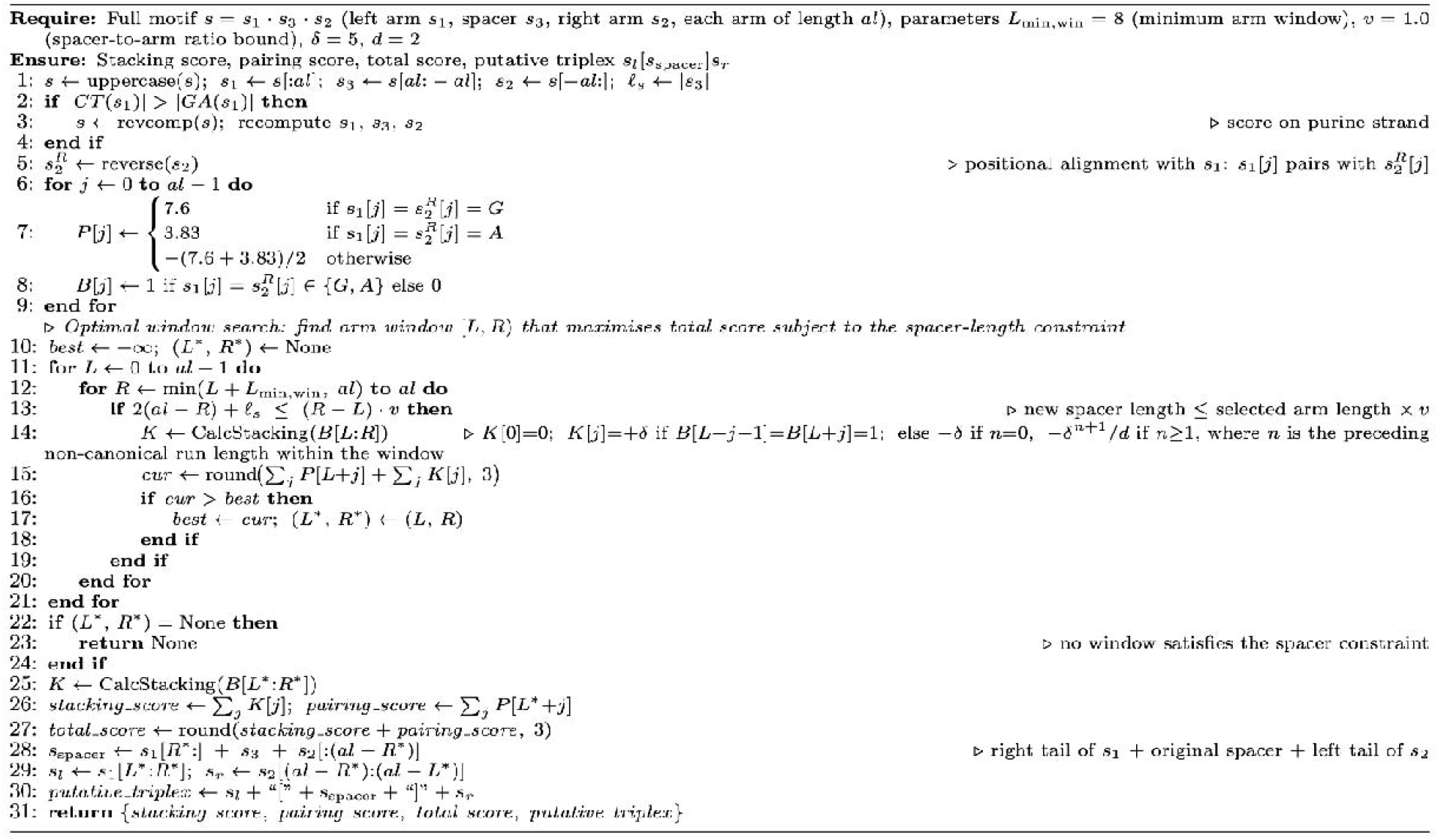

**Algorithm 3: Stability scoring**

Each candidate hit produced by the center-outward detection and overlap removal steps is evaluated for stability. A constrained search over all arm windows (L, R) identifies the contiguous sub-arm that maximises the combined pairing and stacking score, subject to the requirement that the resulting spacer formed by folding the excluded 5′ and 3′ arm tails inward does not exceed the selected arm length. Both arm boundaries are optimised jointly in a single pass, replacing the earlier sequential greedy steps. The result is a stacking score, a pairing score, a combined total score, and a boundary-optimised putative triplex in bracket notation.

## Results

### Computational performance and parallel scalability

To evaluate the computational scalability of HSeeker, we benchmarked the complete analysis pipeline, sequence scanning with the RMQ-based purity prefilter, overlap removal, and stability scoring, on human chromosome 1 (hg38; 248.9 Mbp; FASTA file 252 MB) using a TACC Lonestar6 vm-small virtual machine node. The node provided 16 virtual CPU cores (1 socket, 16 cores per socket, 1 thread per core) of an AMD EPYC 7763 processor operating at 2.45 GHz, with 32 GB of RAM, 256 MB of L3 cache, and running Rocky Linux 8.10 (kernel 4.18.0) with Python 3.11. We varied the number of worker threads from 1 to 16 and recorded wall-clock time and peak resident set size for each configuration, with all other parameters held at their defaults (minimum arm length 8 bp, maximum spacer 10 bp, purity threshold 0.90, RMQ purity prefilter enabled, scoring enabled). All runs produced an identical set of 96,729 non-overlapping H-DNA loci, confirming deterministic output across all parallelisation levels. At a single worker, the full pipeline was completed in 187.7 s (approximately 3.1 min). Increasing the thread count to 2, 4, 8, and 16 yielded wall-clock times of 105.3 s, 59.5 s, 35.9 s, and 25.2 s, corresponding to speedups of 1.78-fold, 3.15-fold, 5.22-fold, and 7.44-fold (**Figure 2**). Parallel efficiency remained above 79% through 4 workers and was 65% at 8 workers and 47% at 16 workers, with the decline at higher thread counts attributable to the post-scan serial phases (cross-chunk overlap removal and stability scoring). Peak memory grew modestly from 1,612 MB at 1 worker to 1,965 MB at 16 workers, consistent with the thread-based design in which additional workers hold more in-flight chunk data within a shared process address space.

**Figure 2:**
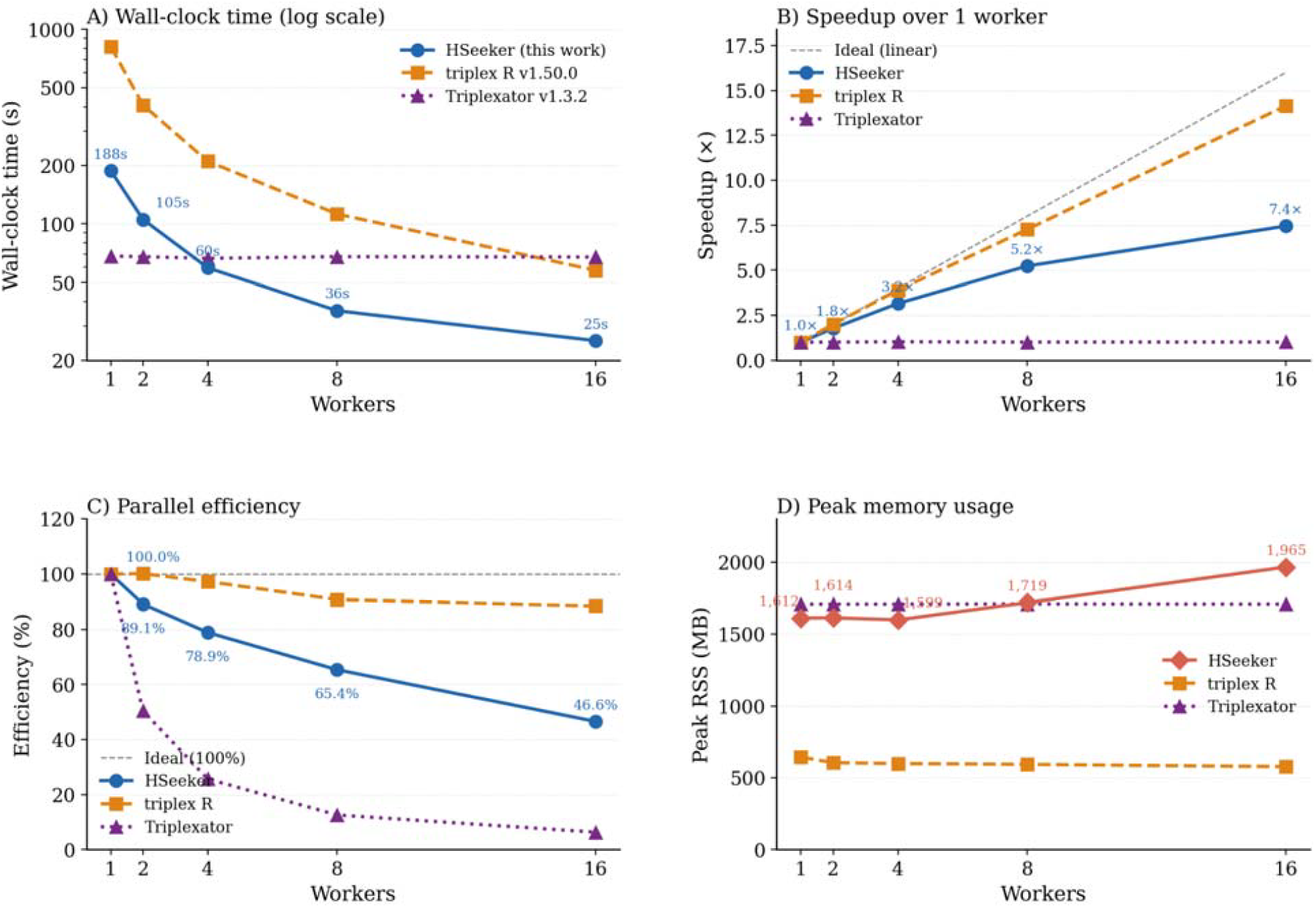
Parallel scaling of HSeeker, triplex R (v1.50.0), and Triplexator (v1.3.2) on human chromosome 1 (hg38, 248.9 Mbp). Wall-clock time (log scale) **A)**, speedup **B)**, parallel efficiency **C)**, and peak memory usage **D)** across 1–16 workers. At 16 workers, HSeeker (25.2 s, 7.44-fold speedup, 47% efficiency) is 2.3-fold and 2.7-fold faster than triplex R (57.7 s) and Triplexator (67.5 s), respectively.

For direct comparison, we benchmarked two established triplex-detection tools, the triplex Bioconductor R package (v1.50.0) and Triplexator (v1.3.2), on the same chromosome and node under sweeps of 1, 2, 4, 8, and 16 workers (**Figure 2**). The triplex package is intrinsically single-threaded; to utilise multiple cores, the chromosome was split into non-overlapping 1 Mbp windows, each processed independently in parallel. This chunk-level decomposition yielded near-ideal scaling, with wall-clock times of 816.3, 407.7, 209.9, 112.4, and 57.7 s at 1 through 16 workers (14.2-fold speedup, 88% parallel efficiency at 16 workers), reporting 81,906 hits. Despite excellent relative scaling, the absolute run time at 16 workers (57.7 s) remained 2.3-fold longer than HSeeker (25.2 s). Triplexator provides native multi-threading support; however, on a single long chromosome, this parallelism strategy yielded wall-clock times of 68.3, 67.8, 66.5, 67.8, and 67.5 s at 1 through 16 threads, essentially flat across all thread counts, indicating that sequential locus enumeration in the duplex pass constitutes the dominant bottleneck with no practical benefit from additional cores. Triplexator reported 275,476 triplex target site loci, reflecting a broader and non-equivalent target class relative to H-DNA mirror repeats; at 16 threads, HSeeker was 2.7-fold faster. Peak RSS was 578 MB for triplex R (low, because each parallel worker holds only a 1 Mbp subsequence) and 1,709 MB for Triplexator (a single process loading the full chromosome), compared with 1,965 MB for HSeeker at 16 workers.

Taken together, these results demonstrate that HSeeker scales efficiently across multiple cores and outperforms both comparators in absolute wall-clock time at every worker count examined. At 16 workers, chromosome 1 is fully analysed in 25 s; by linear extrapolation, the complete human reference genome (approximately 3,200 Mbp) would require approximately 5 min for HSeeker, compared with approximately 12 min for triplex R and approximately 15 min for Triplexator at the same core count, and approximately 2.9 hours for triplex R at a single worker.

### Comparison of HSeeker with other H-DNA detection tools using experimental data

In the sensitivity analysis, we compared HSeeker with the Triplex tool^21^ on a compiled dataset of 79 experimentally validated sequences known to be H-DNA-forming or non-forming (**Supplementary Table 1**). In the primary direct-sequence validation, HSeeker recovered 69 out of 70 experimentally supported H-DNA-forming sequences and rejected 6 of the 9 non-forming controls, corresponding to 98.6% sensitivity and 66.7% specificity (**Supplementary Figure 1**). Triplex^21^ exhibited similar performance in the direct-sequence evaluation with two more false negatives than HSeeker, but improved specificity. On the other hand, Triplexator ^25^ exhibited identical sensitivity but lower specificity than HSeeker. Furthermore, we examined the ability of HSeeker to recover the same dataset randomly inserted in the *E. coli* K-12 genome. HSeeker achieved identical performance, recovering 69 out of 70 H-DNA-forming sequences and rejecting 6 of the 9 non-forming sequences, yielding the same sensitivity and specificity as observed on the original dataset (**Figure 3A**). Triplex^21^ recovered 33 of the 70 H-DNA-forming insertions while rejecting all 9 non-forming insertions (**Figure 3A**). Finally, Triplexator^25^ exposed the same sensitivity as HSeeker but lower specificity by rejecting 5 out of 9 non-forming negative controls. Overall, HSeeker exhibited superior performance (MCC=0.73). Because the injected-genome benchmark depends on genomic context, interval matching, and tool-specific reporting behavior, we treat it as a secondary recovery analysis rather than the primary sequence-level comparison. Additionally, using the HSeeker’s stability-score (**Algorithm 3**), we calculated the ROC curve for various thresholds (**Figure 3B**). Finally, we computed the confusion matrices for both the raw HSeeker output (**Figure 3C**) and the refined confusion matrix (**Figure 3D**), showcasing how the stability score can be used to increase the specificity of HSeeker by assigning a more stringent stability scoring threshold to the body of the mirror repeat sequence.

**Figure 3:**
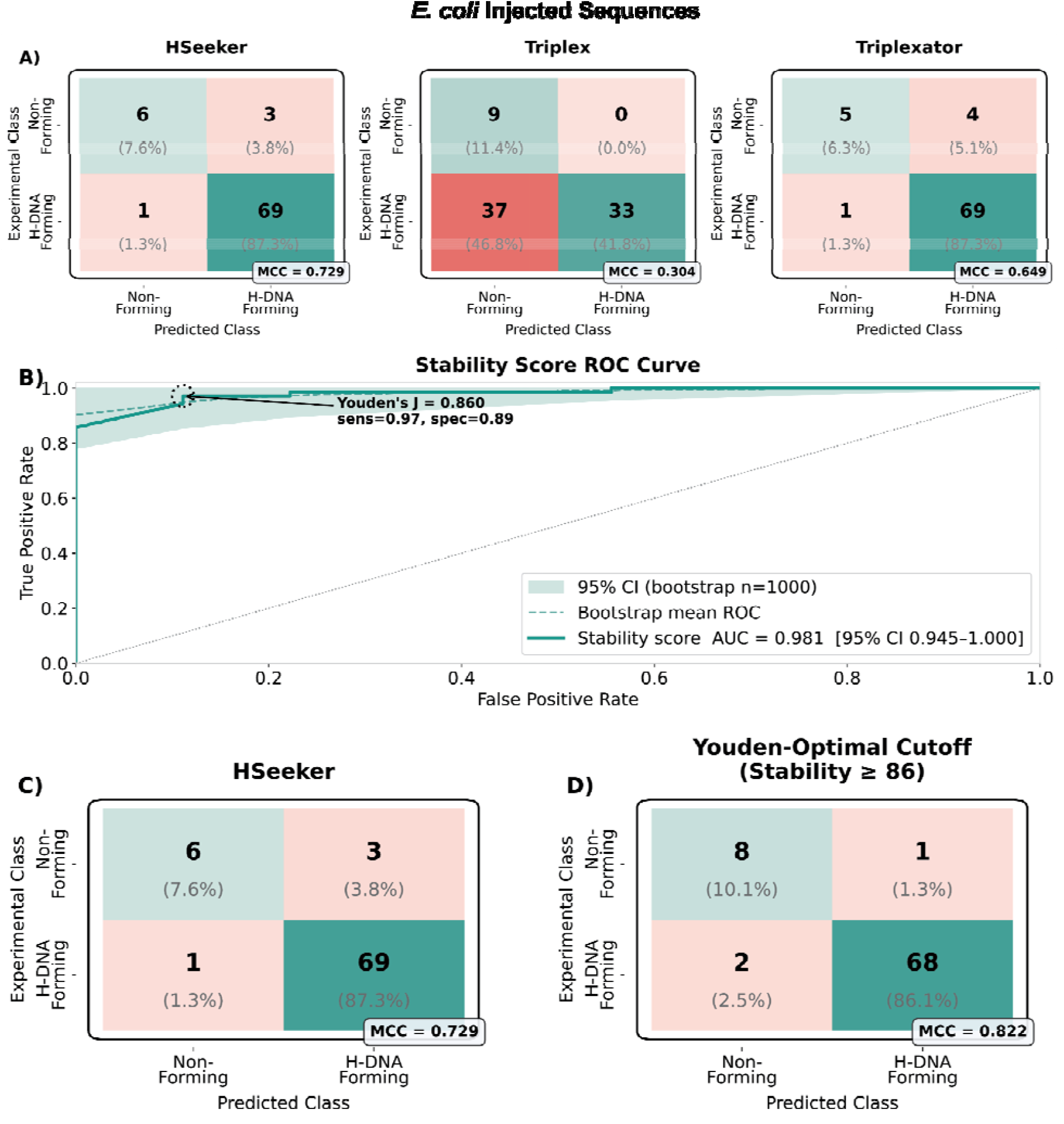
Sensitivity and specificity analysis of HSeeker performance. Experimental H-DNA-forming and non-forming sequences were used to evaluate HSeeker classification performance. 79 curated experimental sequences were randomly inserted into the *E. coli* K-12 reference genome and recovered using an 80% overlap criterion. **A)** Confusion matrices compare experimentally defined sequence classes with predicted classes for HSeeker, Triplex^21^, and Triplexator^25^ recovery of inserted H-DNA-forming and non-forming sequences in genomic context. Confusion matrices compare experimental labels with the tools’ predictions. **B)** Experimental H-DNA-forming (n=70) and non-forming (n=9) sequences were injected into the *E. coli* K-12 genome and used to evaluate HSeeker classification performance. The stability-score ROC curve evaluates the ability of the HSeeker stability score to discriminate H-DNA-forming from non-forming sequences. The 95% confidence interval was estimated using bootstrapping (N=1,000). **C)** Confusion matrix for raw HSeeker detection, where an injected sequence in *E. coli* K-12 was classified as positive if at least one H-DNA candidate was detected. **D)** Confusion matrix after applying the Youden-selected HSeeker stability-score threshold, showing the more specificity-oriented operating point obtained by score-based filtering.

Overall, this validation indicates that HSeeker is at least comparable to Triplex^21^ and Triplexator^25^ on experimentally characterized H-DNA sequences.

### Command-line interface (CLI) tool and package

HSeeker is freely distributed as an open-source package through PyPI and GitHub and can be used either as a command-line interface tool or as an importable Python library. After installation, users can run HSeeker directly from the terminal to analyze FASTA-formatted input sequences, specify detection parameters, and export predicted H-DNA-forming loci for downstream analysis. HSeeker can also be incorporated into custom Python workflows, enabling users to automate analyses, process multiple datasets, or integrate H-DNA detection into larger bioinformatic pipelines. This dual functionality provides flexibility for both standalone use and programmatic applications, supporting users with different levels of computational experience and different analysis needs.

### HSeeker web interface

To improve accessibility and facilitate broader use of HSeeker, we developed a web interface that enables users to identify putative H-DNA-forming sequences without requiring command-line experience. Through the interface, users can submit custom DNA sequences or genomic regions, adjust search parameters, and retrieve annotated candidate H-DNA motifs in a user-friendly format. The web platform provides downloadable results for downstream analysis. By lowering technical barriers to H-DNA detection, the HSeeker web interface enables researchers from diverse backgrounds to incorporate non-B DNA annotation into studies of genome organization, gene regulation, genetic instability, disease etiology, and evolution.

The main detection page allows users to either paste FASTA-formatted sequence data into an input box or upload a FASTA file up to 200 MB (**Figure 4**). The adjustable parameters include minimum and maximum spacer and arm lengths, GA/CT purity threshold in the arms, stability score calculation, and overlap handling. Overlap handling includes either removing overlaps, in which case the sequence with the longest arm is kept, or keeping overlapping H-DNA sequences. Three preset options are also available: i) Strict, ii) Conservative, and iii) Default.

**Figure 4:**
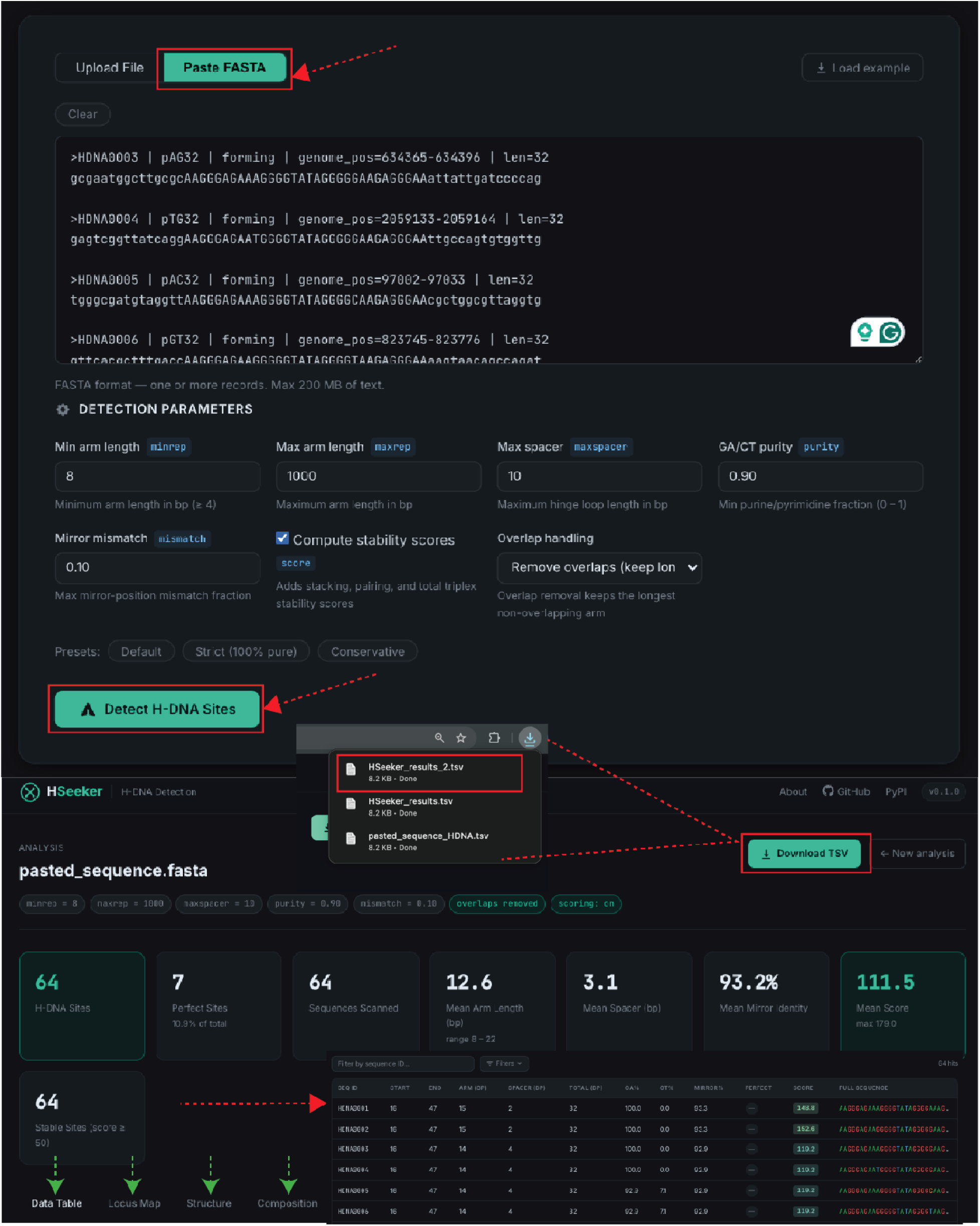
HSeeker web interface. Interface for submitting genomic sequences in FASTA format and configuring job parameters in the HSeeker web app. Detection parameters can be adjusted through the interface, including minimum and maximum arm length, maximum spacer length, GA/CT purity threshold, mirror-mismatch tolerance, stability-score calculation, and overlap handling. After submission, HSeeker reports summary statistics and a full H-DNA sequence table with filtering and dynamic graphs are provided alongside the detection of H-DNA-forming sequences.

The results page is organized into four tabs that provide complementary views of the predicted H-DNA motifs (**Figure 4**). The Data Table tab reports each candidate site in a sortable and filterable table, including sequence ID, start and end coordinates, arm length, spacer length, total motif length, GA/CT composition, mirror identity, perfect-match status, stability score, and full sequence, with the complete table available for download as a TSV file. The Locus Map tab visualizes the distribution of H-DNA sites across input sequences using sequence-level bar plots, a perfect versus imperfect site summary chart, and a genomic-position locus map in which candidate sites are shown as points scaled or colored by arm length. The Structure tab summarizes motif architecture using arm-length and spacer-length distributions, together with an arm-by-spacer density heatmap that highlights common structural configurations among predicted sites. The Composition tab displays sequence-composition and quality-control plots, including GA versus CT composition profiles, mirror-identity distributions, total stability score distributions, and score-versus-mirror-identity scatter plots, enabling users to assess both compositional purity and predicted structural stability (**Figure 5**).

**Figure 5:**
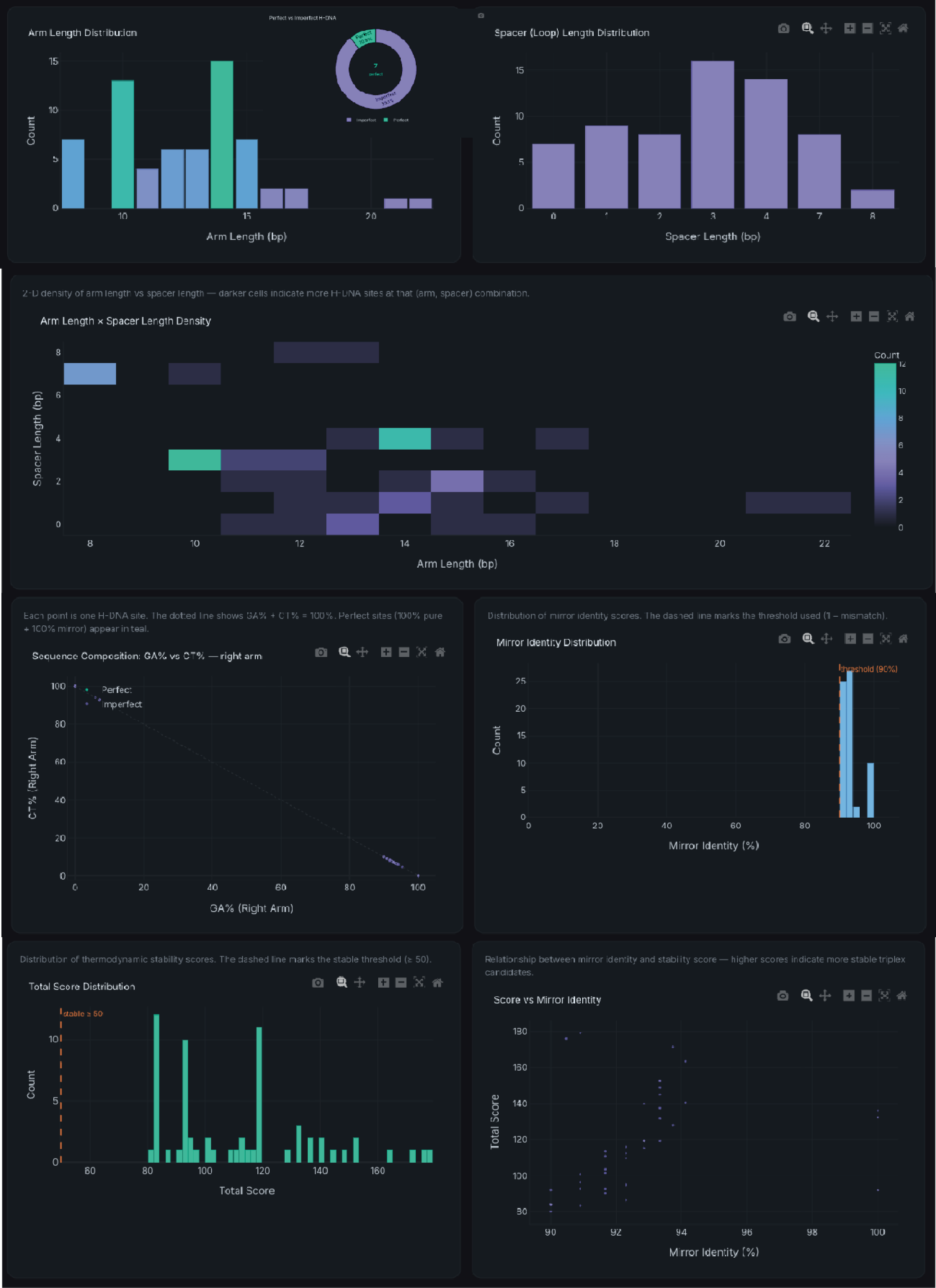
Interactive visualization of predicted H-DNA sequence features in the HSeeker web interface. The interface for H-DNA dynamic graphs enables users to further assess their sequences of interest. An arm-length by spacer-length density heatmap highlights recurrent motif configurations and identifies combinations of arm and spacer lengths that are enriched among the detected candidates. Additional graphs summarize GA-rich and CT-rich arm composition, mirror-identity distributions, total stability-score distributions, and the relationship between stability score and mirror identity.

Finally, we have included an About page, which provides background information on H-DNA biology, the structural requirements for H-DNA formation, the HSeeker center-outward mirror-repeat extension algorithm, output fields, stability scoring, parameter modes, validation, command-line usage, Python API access, and current limitations. Together, these features make the HSeeker website a user-friendly platform for H-DNA sequence detection, visualization, and interpretation.

## Discussion

Here, we present a novel software, HSeeker, for the systematic identification of putative H-DNA-forming sequences. HSeeker addresses an important gap in non-B DNA annotation. HSeeker combines a center-outward mirror-repeat search strategy, greedy overlap removal, and stability scoring to move beyond simple mirror-repeat detection. By integrating experimentally supported physical and thermodynamic features of homopurine triplex formation with biologically relevant determinants of H-DNA stability, including homopyrimidine composition, spacer length, mirror mismatches, arm length, Hoogsteen pairing, base stacking, and boundary optimization, HSeeker generates an overall stability score while maintaining high-throughput performance. Thus, HSeeker provides an interpretable and scalable approach for studying H-DNA-forming potential across genomic datasets.

We also performed thorough benchmarking against two existing H-DNA and mirror-repeat detection tools, Triplex and Triplexator^21,25^, as well as against experimentally characterized H-DNA-forming and non-forming sequences, demonstrating that HSeeker provides improved detection performance and interpretable stability-based prioritization. The list of positive controls was based on rigorous proof of H-DNA formation, such as structure fine-mapping at a nucleotide level via chemical modifications, tested within a plasmid context (**Supplementary Table 1**). Another layer of evaluation involved using S1 and P1 nuclease assays, and H-DNA-induced mutagenesis measured in mammalian cells (for the H-DNA-forming sequence from the *c-MYC* gene)^12^. The Stability Scores predicted by HSeeker agreed with these biochemical and genetic findings^12^. The successful discrimination of experimentally validated sequences confirms that this biophysically informed scoring system works highly effectively and aligns seamlessly with previously published thermodynamic parameters.

3plex^26^ is so far the only other program that has also integrated the biophysical triplex structural features, such as thermal stability information derived from triplex denaturation experiments, into the triplex identification algorithm. But this tool was designed for predicting interactions between single-stranded RNAs and double-stranded DNA regions mediated by intermolecular triple-helix formation, not for intramolecular H-DNA-forming sequences in genomic DNA. HSeeker is the only H-DNA identification algorithm with a proven scoring system built-in, and can achieve high-throughput processing speeds. The center-outward heuristic search algorithm and subsequent greedy overlap removal allow the software to scale linearly with sequence length. HSeeker is optimized for scanning massive inputs with high speed. As demonstrated, it was able to screen the entire human chromosome 1 in approximately 25 seconds using a standard multi-threaded environment. This computational efficiency enables rapid, whole-genome profiling and facilitates broad, cross-species evolutionary comparisons that were previously computationally prohibitive. In addition, HSeeker also provides a user-friendly web interface for easy access, eliminating the technical barriers associated with non-B DNA structural analysis. It allows users to customize the key search parameter, such as the penalty scores for mismatches and the limitations of (Max and Min) spacer and arm lengths between the mirror-symmetric arms, to meet their own project needs.

By systematically mapping these dynamic alternative DNA structures, HSeeker will provide a valuable tool for uncovering the diverse roles of H-DNA in gene regulation and identifying endogenous sources of disease-associated genome instability.

## Code availability

The HSeeker package and its Python bindings are released under the GPL license as a multi-platform application and are available at: https://github.com/Georgakopoulos-Soares-lab/HSeeker. A stable version of HSeeker is also found at: https://pypi.org/project/hseeker/. The HSeeker web interface can be found at https://hseeker-production.up.railway.app/.

## Data Availability

The 79 experimentally validated H-DNA-forming and non-forming sequences used for benchmarking, along with their source references, are provided in **Supplementary Table 1**. The human chromosome 1 sequence (hg38) used for performance benchmarking was obtained from the UCSC Genome Browser (http://hgdownload.soe.ucsc.edu/downloads.html#human). The *E. coli* K-12 reference genome used for genomic-context validation was downloaded from NCBI. All benchmarking results, including sensitivity/specificity analyses, ROC curves, and computational performance data generated in this study, are available from the corresponding authors upon reasonable request and will be deposited in a public repository upon publication.

## Supplementary Figures

**Supplementary Figure 1:**
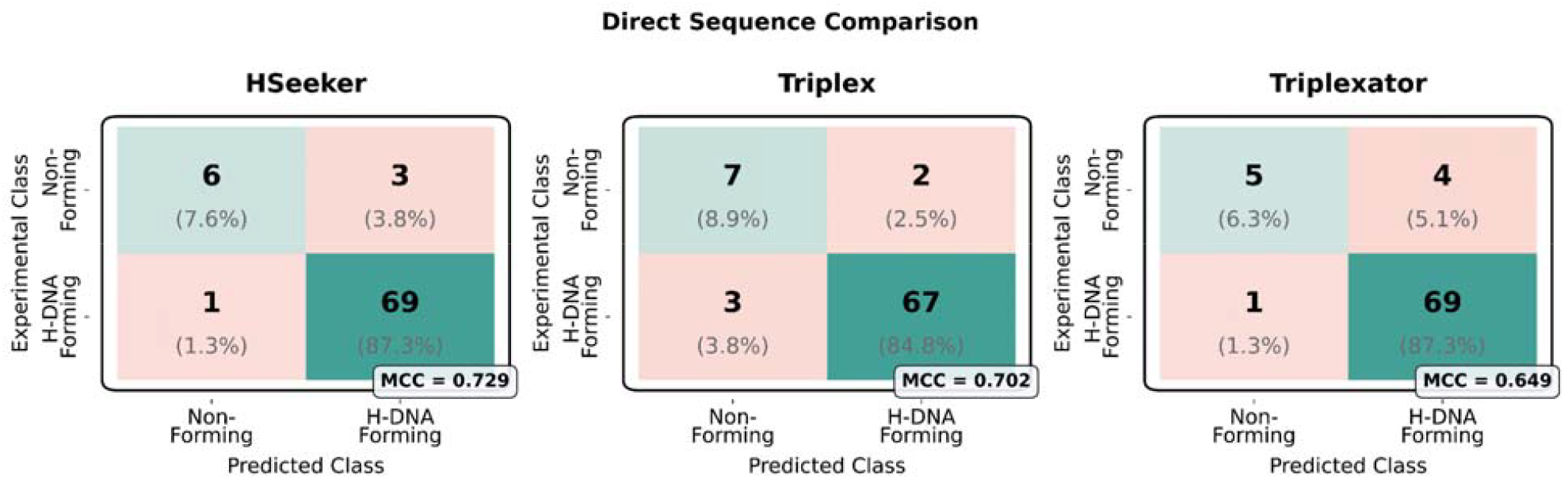
Experimental H-DNA-forming (n=70) and non-forming (n=9) sequences were used to evaluate HSeeker classification performance. Direct sequence-level comparison of HSeeker, Triplex^21^, and Triplexator^25^ on 79 curated experimental sequences. Confusion matrices compare experimentally defined sequence classes with predicted classes for HSeeker, Triplex ^21^, and Triplexator ^25^ recovery of H-DNA-forming and non-forming sequences.

## References

1. Mirkin, S. M. et al. DNA H form requires a homopurine-homopyrimidine mirror repeat. Nature 330, 495–497 (1987).

2. Frank-Kamenetskii, M. D. & Mirkin, S. M. Triplex DNA structures. Annu Rev Biochem 64, 65–95 (1995).

3. Hisey, J. A., Masnovo, C. & Mirkin, S. M. Triplex H-DNA structure: the long and winding road from the discovery to its role in human disease. NAR Mol Med 1, ugae024 (2024).

4. Kouzine, F. et al. Permanganate/S1 Nuclease Footprinting Reveals Non-B DNA Structures with Regulatory Potential across a Mammalian Genome. Cell Syst 4, 344–356.e7 (2017).

5. Jain, A., Wang, G. & Vasquez, K. M. DNA triple helices: biological consequences and therapeutic potential. Biochimie 90, 1117–1130 (2008).

6. Wang, G. & Vasquez, K. M. Impact of alternative DNA structures on DNA damage, DNA repair, and genetic instability. DNA Repair (Amst) 19, 143–151 (2014).

7. Kaushik Tiwari, M., Adaku, N., Peart, N. & Rogers, F. A. Triplex structures induce DNA double strand breaks via replication fork collapse in NER deficient cells. Nucleic Acids Res 44, 7742–7754 (2016).

8. Wang, G. & Vasquez, K. M. Effects of Replication and Transcription on DNA Structure-Related Genetic Instability. Genes (Basel) 8, (2017).

9. Zhao, J., Bacolla, A., Wang, G. & Vasquez, K. M. Non-B DNA structure-induced genetic instability and evolution. Cell Mol Life Sci 67, 43–62 (2010).

10. Wang, G. & Vasquez, K. M. Dynamic alternative DNA structures in biology and disease. Nat Rev Genet 24, 211–234 (2023).

11. Bochalis, E. et al. Non-B DNA structures and their contributions to genetic diversity, aging, and disease. Nucleic Acids Res 54, (2026).

12. Wang, G. & Vasquez, K. M. Naturally occurring H-DNA-forming sequences are mutagenic in mammalian cells. Proc Natl Acad Sci U S A 101, 13448–13453 (2004).

13. Wang, G., Carbajal, S., Vijg, J., DiGiovanni, J. & Vasquez, K. M. DNA structure-induced genomic instability in vivo. J Natl Cancer Inst 100, 1815–1817 (2008).

14. Georgakopoulos-Soares, I. et al. High-throughput characterization of the role of non-B DNA motifs on promoter function. Cell Genom 2, (2022).

15. Bacolla, A., Tainer, J. A., Vasquez, K. M. & Cooper, D. N. Translocation and deletion breakpoints in cancer genomes are associated with potential non-B DNA-forming sequences. Nucleic Acids Res 44, 5673–5688 (2016).

16. Georgakopoulos-Soares, I., Morganella, S., Jain, N., Hemberg, M. & Nik-Zainal, S. Noncanonical secondary structures arising from non-B DNA motifs are determinants of mutagenesis. Genome Res 28, 1264–1271 (2018).

17. Belotserkovskii, B. P. et al. A triplex-forming sequence from the human c-MYC promoter interferes with DNA transcription. J Biol Chem 282, 32433–32441 (2007).

18. Kinniburgh, A. J. A cis-acting transcription element of the c-myc gene can assume an H-DNA conformation. Nucleic Acids Res 17, 7771–7778 (1989).

19. Del Mundo, I. M. A., Zewail-Foote, M., Kerwin, S. M. & Vasquez, K. M. Alternative DNA structure formation in the mutagenic human c-MYC promoter. Nucleic Acids Res 45, 4929–4943 (2017).

20. Schroth, G. P. & Ho, P. S. Occurrence of potential cruciform and H-DNA forming sequences in genomic DNA. Nucleic Acids Res 23, 1977–1983 (1995).

21. Lexa, M., Martínek, T., Burgetová, I., Kopeček, D. & Brázdová, M. A dynamic programming algorithm for identification of triplex-forming sequences. Bioinformatics 27, 2510–2517 (2011).

22. Cer, R. Z. et al. Non-B DB v2.0: a database of predicted non-B DNA-forming motifs and its associated tools. Nucleic Acids Res 41, D94–D100 (2013).

23. Cer, R. Z. et al. Searching for non-B DNA-forming motifs using nBMST (non-B DNA motif search tool). Curr Protoc Hum Genet Chapter 18, Unit 18.7.1–22 (2012).

24. Berselli, M., Lavezzo, E. & Toppo, S. NeSSie: a tool for the identification of approximate DNA sequence symmetries. Bioinformatics 34, 2503–2505 (2018).

25. Buske, F. A., Bauer, D. C., Mattick, J. S. & Bailey, T. L. Triplexator: detecting nucleic acid triple helices in genomic and transcriptomic data. Genome Res 22, 1372–1381 (2012).

26. Masera, M., Cicconetti, C., Ferrero, F., Oliviero, S. & Molineris, I. 3plex Web: An interactive platform for RNA:DNA triplex prediction and analysis. Comput Struct Biotechnol J 27, 3110–3113 (2025).

27. Park, G., Kang, B., Park, S. V., Lee, D. & Oh, S. S. A unified computational view of DNA duplex, triplex, quadruplex and their donor-acceptor interactions. Nucleic Acids Res 49, 4919–4933 (2021).

